# Acoustic and Visual Stimulus Parameters Underlying Sound Symbolic Crossmodal Correspondences

**DOI:** 10.1101/517581

**Authors:** Sara M. List, Kelly McCormick, Simon Lacey, K. Sathian, Lynne C. Nygaard

**Author notes:** Corresponding authors: Lynne C. Nygaard, Department of Psychology, Emory University, College of Arts and Sciences, Atlanta, GA 30322, USA, K. Sathian, Department of Neurology, Milton S. Hershey Medical Center, Penn State College of Medicine, Hershey, PA 17033-0859, USA.

## Abstract

It is often assumed that a fundamental property of language is the arbitrariness of the relationship between sound and meaning. Sound symbolism, which refers to non-arbitrary mapping between the sound of a word and its meaning, contradicts this assumption. Sensitivity to sound symbolism has been studied through crossmodal correspondences (CCs) between auditory pseudowords (e.g. ‘loh-moh’) and visual shapes (e.g. a blob). We used representational similarity analysis to examine the relationships between physical stimulus parameters and perceptual ratings that varied on dimensions of roundedness and pointedness, for a range of auditory pseudowords and visual shapes. We found that perceptual ratings of these stimuli relate to certain physical features of both the visual and auditory domains. Representational dissimilarity matrices (RDMs) of parameters that capture the spatial profile of the visual shapes, such as the simple matching coefficient and Jaccard distance, were significantly correlated with those of the visual ratings. RDMs of certain acoustic parameters of the pseudowords, such as the temporal fast Fourier transform (FFT) and spectral tilt, that reflect spectral composition, as well as shimmer and speech envelope that reflect aspects of amplitude variation over time, were significantly correlated with those of the auditory perceptual ratings. RDMs of the temporal FFT (acoustic) and the simple matching coefficient (visual) were significantly correlated. These findings suggest that sound-symbolic CCs are related to basic properties of auditory and visual stimuli, and thus provide insights into the fundamental nature of sound symbolism and how this might evoke specific impressions of physical meaning in natural language.

## 1. INTRODUCTION

It is commonly held that a fundamental property of language is that it is arbitrary, i.e. the sounds used to describe a referent do not necessarily resemble that referent (de Saussure, 2011, but see Joseph, 2015). Whether language is fundamentally arbitrary or whether “names ought to be given according to a natural process” has been debated since Plato’s *Cratylus* (Ademollo, 2011, p.32). Language does have significant aspects that are non-arbitrary, for example, iconicity, which refers to the resemblance between properties of a linguistic form and the sensorimotor and/or affective properties of referents (Perniss & Vigliocco, 2014). One type of iconicity is sound symbolism, which includes a broad set of phenomena that show a clear resemblance between properties of speech sounds and properties of their referents, such as onomatopoeia or Japanese mimetics (e.g. ‘gorogoro’ meaning the sound of a heavy object rolling around: Imai & Kita, 2014).

Because sound symbolism embraces a broad set of non-arbitrary phenomena, it has proven difficult to study systematically, but sound-symbolic crossmodal correspondences (CCs) have provided researchers with a unique framework with which to study sound symbolism. A well-known example of sound-symbolic CCs was first described by Köhler (1929) in which individuals displayed a correspondence between a curvy, cloud-like shape and the pseudoword “baluma” and between an angular star-like shape and the pseudoword “takete”.

Sound-symbolic CCs constitute one type of CC, i.e., near-universally experienced associations between seemingly arbitrary stimulus features in different senses (Spence, 2011). CCs often occur between stimulus properties that are correlated (i.e. small animals often emit higher-pitched noises than large animals), which can lead to more efficient processing of sensory information. For example, large and small size in the visual domain are consistently associated with low- and high-pitched sounds (Gallace & Spence, 2006), and high and low elevation of visual stimuli are associated with high and low pitch in the auditory domain (Ben-Artzi & Marks, 1995).

Sound symbolism has been demonstrated across different languages (Blasi et al., 2016), cultures (Chen et al., 2016; Kantartzis et al., 2011), and even with children of pre-reading age (Imai et al., 2015; Maurer et al, 2006; Ozturk et al., 2013; Tzeng, Nygaard, & Namy, 2017). These studies show that the existence of sound-symbolic CCs in language is both prolific and robust. Further, language users are sensitive to sound-symbolic correspondences in that they actively utilize these associations to correctly assign meaning to foreign synonym-antonym pairs at above-chance levels (Nygaard, et al., 2009; Revill et al., 2014; Tzeng, Nygaard, & Namy, 2016). Sound-symbolic CCs also seem to play a role in language processing and early word learning (Imai & Kita, 2014). For example, individuals exhibit sensitivity to sound-symbolic crossmodal associations as early as four months of age (Ozturk et al., 2013), and recent studies have suggested that sound symbolism is important for specific word-to-meaning associations in young children with limited vocabularies (Gasser, 2004; Tzeng, Nygaard, & Namy, 2017). In adults, sound symbolism may offer linguistic processing advantages for categorization and word learning, particularly for larger vocabularies (Brand et al., 2018; Gasser, 2004; Revill et al., 2018). Though most of the evidence supporting sound symbolism has consisted of behavioral studies, more recently neuroimaging studies have begun to reveal the neural correlates of this phenomenon (Revill et al., 2014; McCormick et al., 2018; Peiffer-Smadja & Cohen, 2019).

Two open questions on sound-symbolic CCs are which physical features in the auditory domain come to be associated with which features in the visual domain; and how these features combine to drive a sound-symbolic association. Previous research has suggested that certain phonetic features are more likely to elicit sound-symbolic associations (Nielsen & Rendall, 2011; Fort, Martin, & Peperkamp, 2014; McCormick et al., 2015). For example, obstruents, including the English consonant sounds ‘p’, ‘t’, and ‘k’, are more likely to evoke pointedness, and sonorants, including the English consonant sounds ‘l’ and ‘m’, are more likely to evoke roundedness (McCormick et al., 2015). In addition, the relevant phonetic features can change depending on the visual association: Knoeferle et al. (2017) found that the first formant predicted visual size judgments, the third formant predicted visual shape judgments, and both size and shape judgments were predicted by the second formant. However, the acoustic and visual drivers of sound symbolism (e.g. the spectrotemporal features and amplitude variations of speech sounds and the spatial features of shapes) have received limited examination. In tasks that test artificial language learning, non-arbitrary word-meaning associations are easier to learn than arbitrarily constructed word-meaning associations (Brand et al., 2018; Nielsen, 2016; Revill et al., 2018), but researchers do not know what stimulus features participants attend to during the perception of these artificial words. Assessing the physical features that contribute to a sound-symbolic CC, as we do here, provides insight into the fundamental perceptual processes that drive word-to-meaning associations.

In the experiments reported here, we used sets of systematically varied auditory pseudowords and visual shapes, each of which was rated by participants along dimensions of roundedness and pointedness. Similarities among these perceptual judgments were compared between modalities, and to similarities among acoustic and visual parameters of the stimuli. In order to compare across auditory and visual modalities and between perceptual ratings and various parameters of the stimuli, we used representational similarity analysis (RSA; Kriegeskorte et al., 2008). This method relies on generating representational dissimilarity matrices (RDMs) and has been applied to capture the neural dissimilarity between individual stimuli or classes of stimuli (Kriegeskorte et al., 2008). Here we use RSA in a novel context by comparing the patterns of dissimilarity between judgments of systematically varied shape and sound stimuli and measurements of acoustic and visual parameters of these stimuli. RSA allows us to develop a comprehensive view of the commonalities and the differences in the dissimilarity patterns that relate to both visual and auditory stimuli in the perception of sound-symbolic CCs. The present study was designed to systematically determine which acoustic features best characterize sound-symbolic stimuli and which visual features are related to judgments of shape, using RSA. Specifically, we examined whether a key set of acoustic and visual parameters reliably predicted judgments of roundedness and pointedness of the sounds of pseudowords and the shapes of two-dimensional objects.

## 2. MATERIALS AND METHODS

### 2.1 Participants

A total of 61 Emory University students (28 male, 33 female, mean (± standard deviation (SD)) age 20 ± 4 years) gave informed consent and received course credit for their participation. Thirty participated in a Likert-type rating task for visual shapes (14 male, 16 female) and a separate 31 participated in a Likert-type rating task for auditory pseudowords (14 male, 17 female). All participants were native English (American) speakers and reported normal or corrected-to-normal vision and no known hearing, speech, or language disorders. All procedures were approved by the Emory University Institutional Review Board.

### 2.2 Stimuli

#### 2.2.1 Shapes

Ninety shapes were created using Adobe Illustrator (Ventura, CA) and consisted of gray line drawings (RGB: 240, 240, 240) on a black background. Shapes had four, five, or six protuberances. Shapes were created using a template of three concentric circles (25, 35, and 45 mm radii), the outer circle serving as a bounding border with protuberances extending to its perimeter. The two inner circles served to define the inward extent of each protuberance. Thinner protuberances (30 shapes) extended all the way to the innermost circle; thicker protuberances (30 shapes) extended only to the middle circle; the remaining 30 shapes were constructed with a mix between thin and thick protuberances. For each shape of one category (rounded or pointed), there was a corresponding shape in the other category with the same outer and inner anchor points, which created fifteen thick, fifteen thin, and fifteen heterogeneous shapes in each category. Under these constraints, the full complement of shapes included a rounded and pointed shape with the same inner and outer anchor points.

#### 2.2.2 Pseudowords

570 pseudowords were constructed using only phonemes and combinations of phonemes that occur in the English language (McCormick et al., 2015). Each pseudoword consisted of two syllables of the form: consonant, vowel, consonant, vowel (CVCV). Consonants were sampled from sonorants, fricatives/affricates, and stops; of the obstruents, including fricatives/affricatives and stops, half were voiced and half were unvoiced. Vowels were either front/rounded or back/unrounded. For a complete description of the stimulus set, see McCormick et al. (2015).

The pseudowords were recorded by a female native speaker of American English with neutral intonation. Pseudowords were read from a computer screen and recorded in a sound-attenuated room using a Zoom 2 Cardioid microphone and digitized directly onto a Dell computer operating Windows Vista with an Intel Core2 processor at a 44.1 kHz sampling rate. Each pseudoword was then down-sampled at 22.05 kHz (standard for speech and to allow for consistent sample sizes in acoustic measurements of the pseudowords) and amplitude-normalized using PRAAT (Boersma & Weenink, 2012), a speech analysis software package. The pseudowords had a mean (± SD) duration of 457 ± 62 ms.

Each pseudoword was checked by four raters for accurate pronunciation of the intended segments and mispronounced pseudowords were re-recorded and edited, after which they were re-checked by the same four raters. Thirty-three of the pseudowords were rated by at least one of four independent listeners to be real English words and were therefore excluded, for a final total of 537 pseudowords (McCormick et al., 2015).

### 2.3 Likert-type rating tasks

The 537 pseudowords were presented auditorily, once each, over Beyerdynamic DT100 headphones at approximately 75db SPL for one set of participants (n = 31). For a separate set of participants (n = 30), the 90 shapes were presented sequentially, once each, at the center of a Windows 7.0 desktop computer screen using EPrime software Version 2.0.8.22 (Schneider, Eschmann, & Zuccolotto, 2002). The stimuli were presented in pseudorandom order.

Participants were randomly assigned to one of four Likert-type ratings tasks in which they were asked to rate pseudowords or shapes using one of two 7-point Likert-type scales: the first rated pointedness from 1 (not pointed) to 7 (very pointed) and the second rated roundedness from 1 (not rounded) to 7 (very rounded). For pseudowords, 16 participants used the pointedness scale and 15 the roundedness scale (total n = 31). To discourage participants from matching pseudowords with a specific word in the instructions (e.g. ‘peh-kee’ and ‘pointed’), the instructions included several related terms for the concepts of rounded and pointed. For the shapes, 13 participants used the pointedness scale and 17 the roundedness scale (total n = 30). For both pseudowords and shapes, the 7-point scale appeared on the screen on each trial, either in the center of the screen for pseudowords or below each shape. The response keyboard always had 1-7 listed from left to right.

### 2.4 Data Processing and Analysis

#### 2.4.1 Behavioral Analysis

Behavioral data were analyzed in MATLAB 2016a (The MathWorks, Natick, 2016), visual parameters were analyzed using MATLAB, and acoustic parameters were analyzed using both MATLAB and PRAAT (Boersma & Weenink, 2012). We implemented RSA using custom-built code in MATLAB to analyze how each of our acoustic and visual parameters of interest related to the perceptual RDMs. This method involves computing second-order correlations, between a reference RDM and test RDMs (Figure 1).

**Figure 1.**
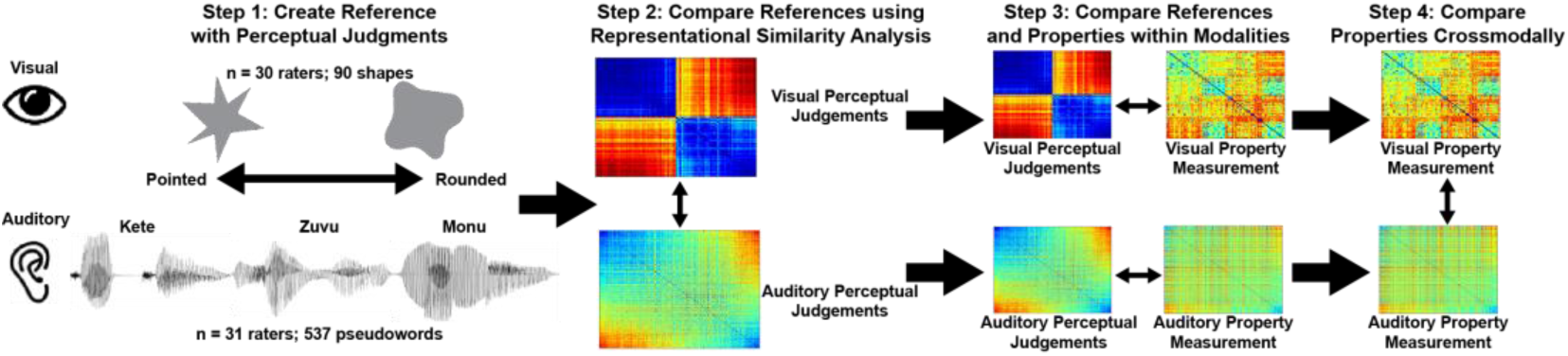
Analysis pipeline. In step 1, we created reference RDMs based on the perceptual judgments in the Likert task. In step 2, we compared the perceptual judgments of the shapes to those of the pseudowords. In step 3, we then compared RDMs of perceptual judgments to RDMs of various measurements of the stimuli for both shapes and pseudowords. In step 4, we compared RDMs of selected visual parameters of the shapes to those of selected auditory parameters of the pseudowords.

To first create a reference RDM based on the perceptual judgments (Figure 1, step 1), the pseudowords were ordered from most pointed to most rounded based on the mean Likert rating of each stimulus. When ordering the stimuli for the perceptual RDM models, the two Likert scales (1-7: not pointed to very pointed, and 1-7: not rounded to very rounded) were recoded such that 1 was equal to “not pointed” on one scale and “very rounded” on the other, and 7 was equal to “very pointed” on the first scale and “not rounded” on the second. However, once the stimuli were thus ordered, we used the original (un-recoded) data within the perceptual RDMs because the RDMs rely on variation, and we are here interested in comparing across pseudowords or across shapes, rather than comparing across participants: regardless of the scale that any individual participant used to rate the stimuli, we are interested in how the patterns of the judgments were dissimilar across the pseudowords or shapes. As illustrated by Figure 2, the judgments for the pseudowords and for the shapes were comparable across the two Likert scales.

**Figure 2.**
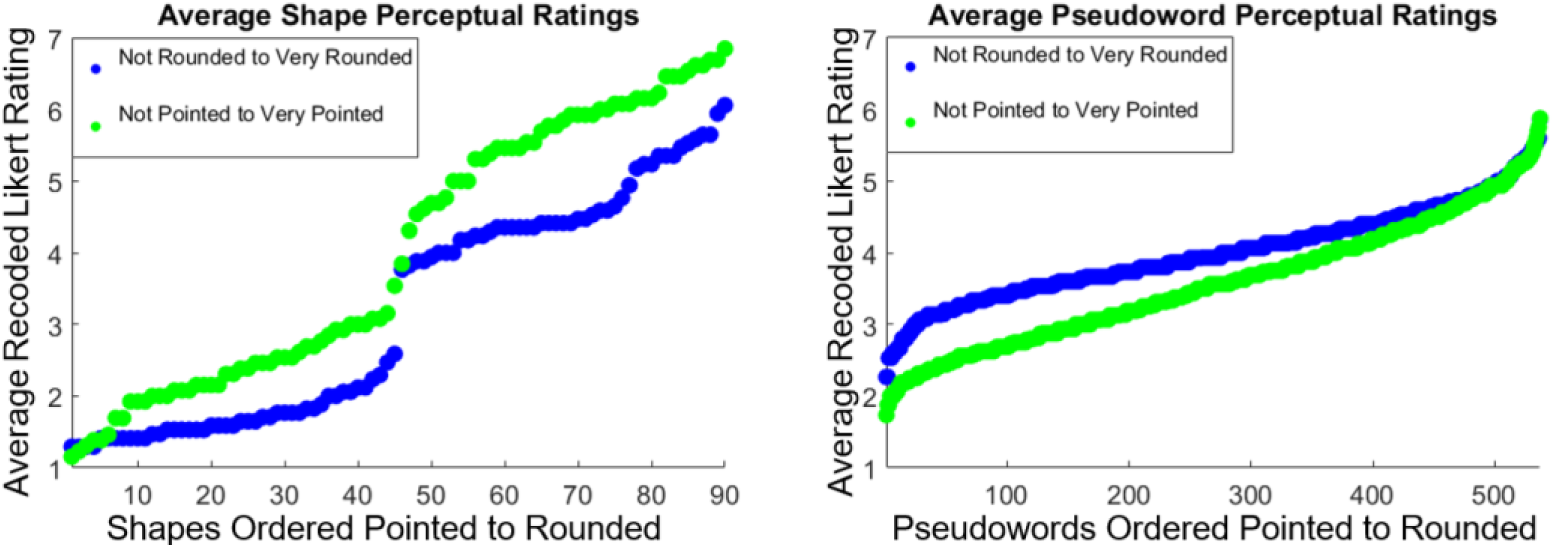
The recoded average rating for each shape (left) and pseudoword (right) stimulus across the two Likert scales.

First, we compared the perceptual judgments RDMs across modalities (Figure 1, step 2). The RDMs represent the pairwise dissimilarity between stimuli, given by 1-Pearson’s correlation coefficient. Then we compared auditory and visual perceptual RDMs to their respective within-modal property RDMs (Figure 1, step 3). Using all participant ratings for shapes and for pseudowords respectively, we constructed two RDMs: one of 90 by 90 for shapes and the other of 537 by 537 for pseudowords. Each of these RDMs was used to compare with RDMs for specific parameters of the corresponding modality (see below).

For the pseudowords, when comparing perceptual judgments to parameters that comprised a single value per pseudoword using second-order correlations (see Table 4), we down-sampled the RDM for the perceptual judgments and binned the acoustic measurements to create 18×18 matrices with 30 pseudowords within each cell. This provided a sufficient number of measures to compute a correlation coefficient that could be entered into each cell of an RDM, while still allowing a sizable enough matrix to be compared with another RDM.

Finally, we compared significant shape and pseudoword parameters across modality (Figure 1, step 4). To compare across modalities, we down-sampled the RDM for pseudowords, which was already ordered via the perceptual ratings, by selecting every sixth pseudoword from the original 537 by 537 matrix to create a 90 by 90 matrix for pseudowords that could then be compared with the 90 by 90 matrix for shapes. To compare RDMs, we used Spearman’s r for second-order correlations instead of Pearson correlations because Spearman’s r is a non-parametric measurement that does not rely on the assumption of normality of the data.

#### 2.4.2 Analysis of stimulus parameters

After analyzing the behavioral results and creating the perceptual RDMs, several visual and acoustic parameters of the stimuli were measured and used to generate RDMs for each parameter (Figure 1, step 3). To test the relationship of the different stimulus parameters to the perceptual ratings in each modality, second-order Spearman correlations between the RDMs for each parameter and the perceptual ratings in that modality were computed. Note that not all stimulus parameters required the same number of samples per image or per pseudoword, respectively, which may serve important when comparing effect sizes in the results (Tables 1 and 2).

**Table 1.** Description of visual parameters of shapes that were compared to the reference RDM from the perceptual judgments.

| Visual Parameters | Description |
| --- | --- |
| Jaccard Distance | The Jaccard distance has been used for RSA of images (Devereux et al., 2013) and is defined as one minus the size of the intersection of the shapes divided by the size of the union of two sample shapes. The Jaccard distance, a variation on the simple matching coefficient, more specifically is $1 - (a)/(a+b+c)$, where the components include the number of pixels present (a) in both images, (b) in the first image but not in the second, (c) in the second image but not in the first (Ricotta & Pavoine, 2015). This pairwise measure is included in the RDM for this parameter as the Jaccard distance instead of 1- Pearson r (160,000 samples/image). |
| Simple Matching Coefficient | The simple matching coefficient is an example of the use of presence/absence coefficients based on the matching and mismatching pixels within two images. The components of these coefficients include the number of pixels present (a) in both images, (b) in the first image but not in the second, (c) in the second image but not in the first, and (d) the pixels absent from both images but found in other images, such that the sum $S = a + b + c + d$ is the total number of pixels in the entire collection of stimuli (Ricotta & Pavoine, 2015). The simple matching coefficient is given by $(a+d)/(a+b+c+d)$. This pairwise measure is included in the RDM for this parameter as $1 - \text{simple matching coefficient}$ instead of $1 - \text{Pearson } r$ (160,000 samples/image). |
| Spatial Frequency FFT | The spatial frequency FFT captures how often the components from the FFT repeat within an image per unit distance (cycles per millimeter; 40,000 samples/image). |
| Silhouette | Each of the shapes was converted into a series of zeros (white) and ones (black) such that each grey shape was converted to white so that it appeared as a silhouette against a black background. The shapes were then reduced from a two-dimensional shape into a single one-dimensional vector, and the RDM was constructed as for image outlines (see below; 40,000 samples/image). |
| Binary Images | Each of the shapes was converted into a series of zeros (white) and ones (black) such that both the outline (white) and silhouette (black) were included. The shapes were then reduced from a two-dimensional shape into a single one-dimensional vector, and the RDM was constructed as for image outlines (see below; 160,000 samples/image). |
| Image outlines | Each of the shapes was converted into a series of zeros (white) and ones (black) in which only the outline of each shape was included. The shapes were then reduced from a two-dimensional shape into a single one-dimensional vector. The resulting vectors were used to compute $1 - \text{Pearson } r$ for the RDM (160,000 samples/image). |

**Table 2.**
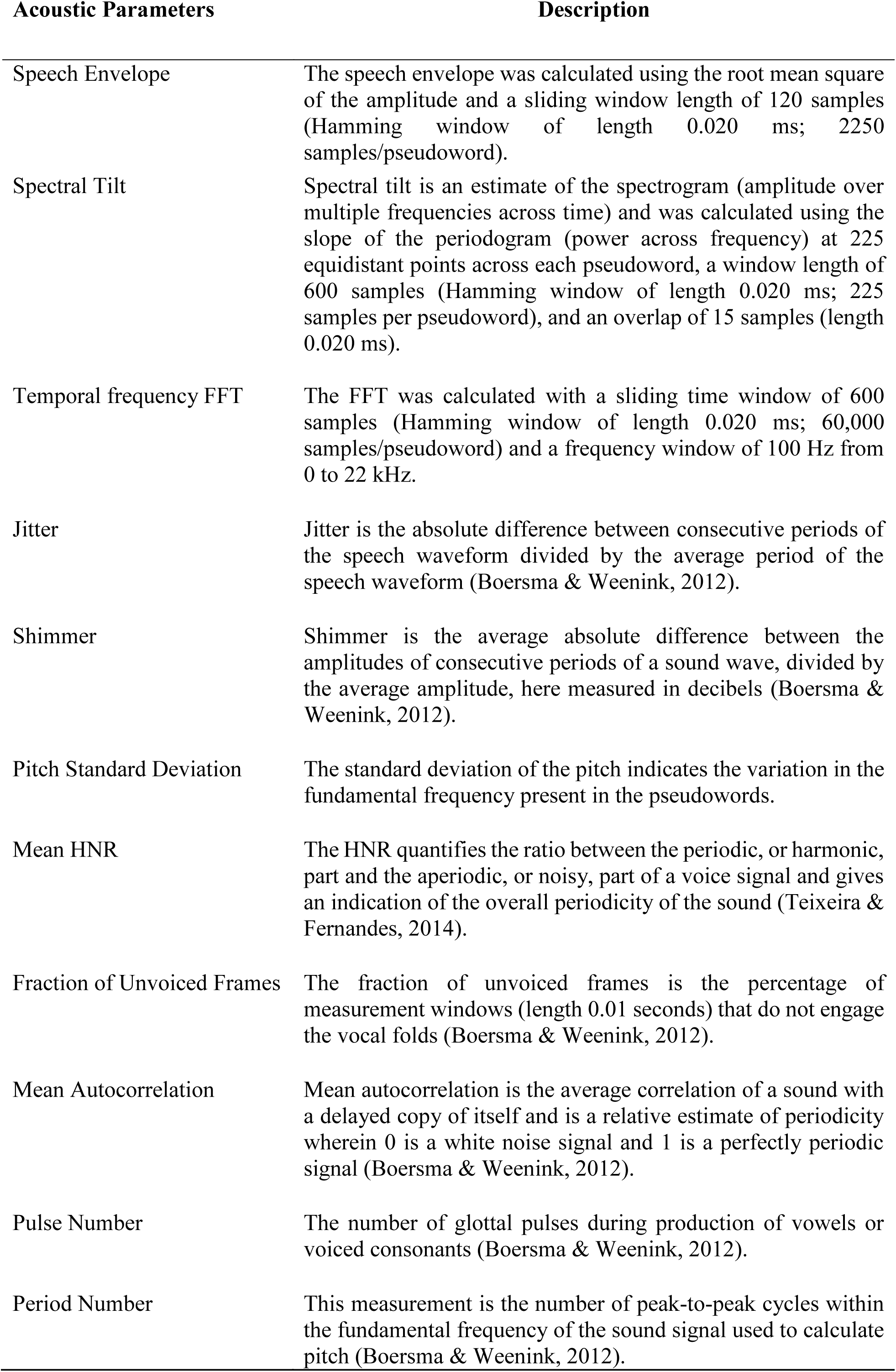
Description of acoustic parameters of the pseudowords that were compared to the reference RDM from the perceptual judgments.

We also compared selected stimulus parameters across modalities (Figure 1, step 4) in order to determine which were most likely to be driving sound-symbolic CCs. Once we determined which stimulus parameters had RDMs that were most correlated with those of the perceptual judgments, we used the RDMs of the two most highly correlated parameters from the visual domain to compare with highly correlated parameters from the auditory domain. We chose the two most highly correlated parameters in the auditory domain that would still allow us to maintain a consistent matrix size of 90 by 90 across modalities. We then compared RDMs of the two stimulus parameters in each modality with the two in the other modality.

##### Shapes

The visual parameters examined for the shapes were the Jaccard distance, the simple matching coefficient, and the fast Fourier transform (FFT) in the spatial frequency domain, all of which index spatial differences among the shapes, together with the silhouettes, binary images and image outlines, all of which attempt to account for shape in a pixel-wise manner (see Table 1 for computational details).

The Jaccard distance and simple matching coefficient (the latter is also known as the Rand similarity coefficient) are two types of distance measures that have been used to compare similarity and diversity, including patterns in DNA (Deagle et al., 2017), images (Devereux et al., 2013), and voices (Cernak et al., 2016). The spatial frequency FFT has been used to filter noisy or complex patterns in complex image sets (Petrou & Petrou, 2011). The spatial FFT has also been used to access the geometric characteristics of complex images, such as determining the spatial characteristics of retinal microglia (Zhang et al., 2016). The silhouettes of images have been shown to predict activation patterns in early visual cortex (Kriegeskorte et al., 2008) and have been used to reveal overlap in the neural processing of words with the processing of objects (Devereux et al., 2013). Binary images and image outlines capture the entire shape in the form of a vector but may be prone to noisiness that does not capture the spatial patterns among different shapes (Devereux et al., 2013). For all visual parameters, shapes were first rotated such that they aligned as closely as possible by co-registering the points or the center of the curves of each shape with one another.

##### Pseudowords

The auditory parameters examined for the pseudowords (see Table 2 for computational details) were the speech envelope, spectral tilt, and FFT (here in the temporal frequency domain) all of which capture the complexity of the sound structure of each pseudoword across multiple measures of time or of frequency. By contrast, jitter, shimmer, pitch standard deviation, mean HNR, the fraction of unvoiced frames, mean autocorrelation, pulse number, and period number are all measurements that capture a single value for each pseudoword that represents an average or overall measurement of different parameters of the speech signal. For example, jitter and shimmer capture average variations in the glottal waveform, the standard deviation of the pitch captures average variation in the fundamental frequency, and mean HNR captures the average relative periodicity of the speech waveform for a given pseudoword. Even these parameters that arguably capture less complexity because they only provide a single measurement could give insight into the parameters that drive sound-symbolic judgments in the auditory domain. For example, the fraction of unvoiced frames is a measure of voicing, which was found to be important in our phonetic analysis of these pseudowords such that unvoiced pseudowords were more likely to be perceived as pointed (McCormick et al., 2015).

The speech envelope is a measure of the amplitude profile across time and captures energy changes corresponding to phonemic and syllabic transitions (Aiken & Picton, 2008). The speech envelope can visually capture the ‘shape’ of the pseudowords (see Figure 1, step 1, for examples), and is a clear candidate for driving sound symbolic CCs. Spectral tilt contributes to the intelligibility of a speech signal under noisy conditions (Lu & Cooke, 2009). The temporal FFT reflects the power spectrum of the component frequencies and was measured at successive time points in the speech signal to capture the change in frequency composition across time (Singh, 2015). Jitter can be used to identify voice pathologies and stems from variation in the vibration of the vocal cords (Teixeira & Fernandes, 2014), giving rise to instability of sound frequency. Higher shimmer is correlated with the presence of noise and breathiness in the voice (Boersma & Weenink, 2012; Teixeira & Fernandes, 2014) and results from instability of sound amplitude. The standard deviation of the pitch or fundamental frequency has been implicated as an indicator of emotional and mental state, indicating that this measure can acoustically convey meanings that are not limited to the linguistic content (Kliper et al., 2015). The HNR gives an indication of the overall periodicity of a sound and has been used as an index of vocal aging or hoarseness (Ferrand, 2002). Mean autocorrelation is also a relative measure of periodicity of a signal. Since our stimuli were spoken by the same speaker taking care to keep prosody consistent, fundamental frequency should have remained relatively consistent, but pulse number may indicate variation in periodicity or phonation across stimuli (Hollien et al., 1977). All pseudowords were first normalized to a duration of 500 ms. Except where noted for the parameters of speech envelope, spectral tilt, and temporal FFT, RDMs comprised of a single sample or measurement per pseudoword.

## 3. RESULTS

### 3.1 Comparison of visual and auditory perceptual ratings

First, we compared visual and auditory perceptual ratings across modalities (Figure 1, step 2). The pattern of judgments evident in the 90×90 RDM for the pointed/roundedness ratings for the visual shapes (Figure 3, left) was essentially binary. This pattern indicates that participants rated shapes largely as more pointed or more rounded, with only a few shapes considered intermediate. Figure 3 (right) depicts the sub-sampled 90×90 RDM of pointed/roundedness ratings for the pseudowords, which appeared to be more graded. Yet, the two matrices for the shapes and pseudowords were positively correlated (Spearman r = 0.66, p<0.0001). This finding indicates that, even though independent sets of participants judged shapes and pseudowords, the judgments were crossmodally consistent. One may expect this result, given that participants were asked to rate stimuli along dimensions of pointedness or roundedness in both modalities, but for the large set of pseudowords, consistent ratings using a scale that is primarily defined by vision was not guaranteed. The strong positive correlation between the auditory and visual RDMs establishes the presence of sound-symbolic CCs between a large set of 90 auditory pseudowords and 90 visual shapes. This lays the foundation for our subsequent analyses of the stimulus parameters relevant to the CC.

**Figure 3.**
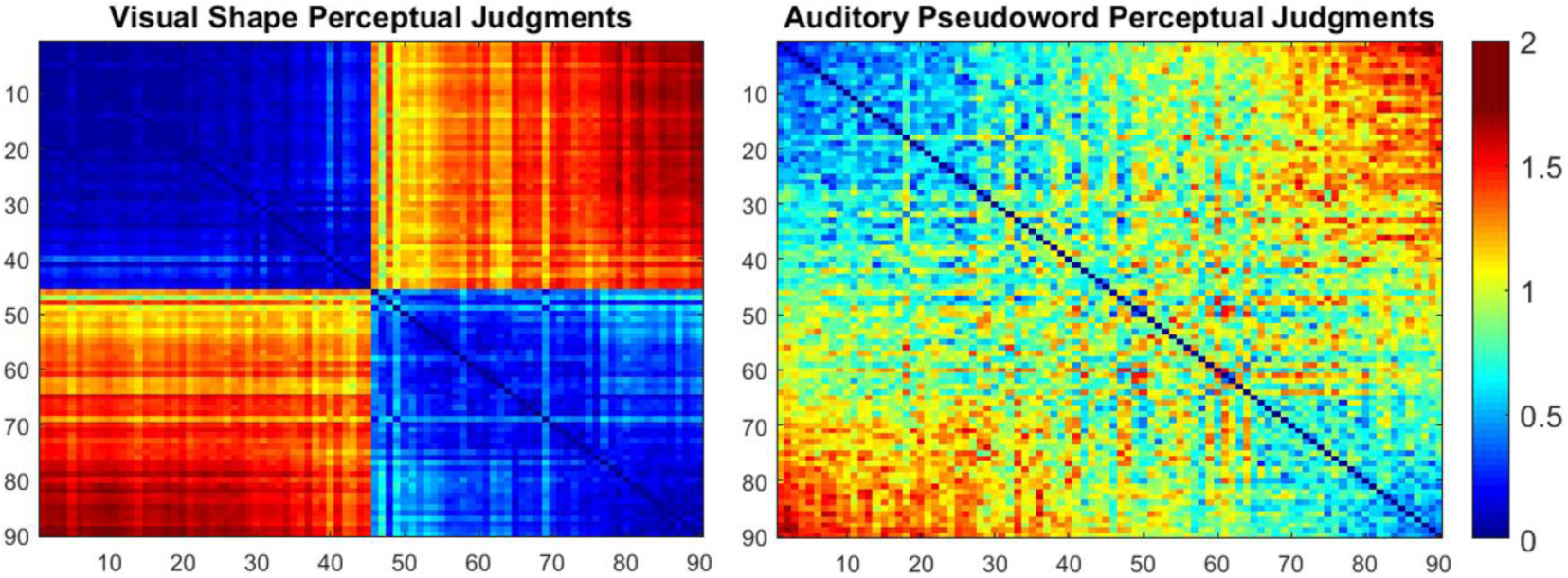
RDMs ordered according to mean perceptual judgments for visual shapes (left) and auditory pseudowords (right) from 1 (rated most pointed) - 90 (rated most rounded). Color bar shows dissimilarity increasing from 0 - 2, which applies to all figures with RDMs. Based on the pattern of correlations, visual shape judgments tended to be binary. Although some pseudowords were judged to be intermediate, there was a significant positive correlation between the two RDMs (Spearman r = 0.66, p < 0.0001).

### 3.2 Comparison of visual perceptual ratings to visual shape indices

Then we compared the visual perceptual RDM to its within-modal visual parameter RDMs (Figure 1, step 3). To determine what properties may influence the judgment of visual roundedness or pointedness, we next compared the RDM for visual judgments to each of the RDMs for the quantitative indices of shape. As an example, Figure 4 compares the dissimilarity matrix of pointedness/roundedness ratings for the shapes (left) to that for the simple matching coefficient (right). These two matrices were positively correlated (Spearman r = 0.28, p<0.0001), although the perceptual judgment RDM was clearly binary in comparison to that for the simple matching coefficient.

**Figure 4.**
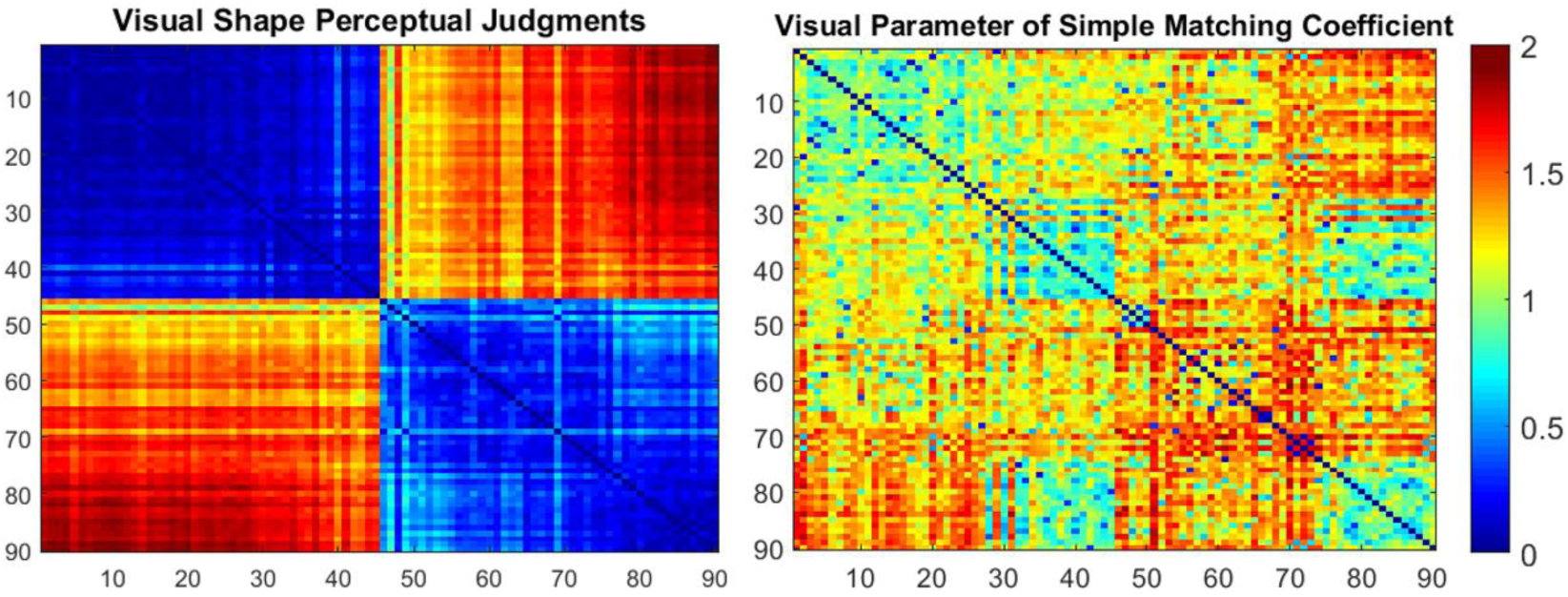
RDMs for the perceptual judgments for visual shapes (left) and one index of shape, the simple matching coefficient (right). Both RDMs are ordered according to mean perceptual judgments for shapes from 1 (rated most pointed) - 90 (rated most rounded). There was a positive correlation between the two RDMs, indicating that the simple matching coefficient is related to visual perceptual judgments of pointedness/roundedness (Spearman r = 0.28, p < 0.0001).

Table 3 shows that the RDM for shape judgments was positively correlated not only with that for the simple matching coefficient as noted above, but also with the RDMs for the silhouette, and Jaccard distance, indicating that these indices of the shapes represent parameters that are important for the visual judgment of roundedness or pointedness. Note that not all of the visual indices required the same number of samples per image, which may affect the comparison of effect sizes (Methods, Table 1). The RDMs for binary images, spatial FFT, and image outline were not significantly correlated with the RDM for shape judgments, indicating that these parameters are less important for the judgment of visual rounded or pointedness.

**Table 3.** Correlations between RDMs for visual perceptual judgments and quantitative parameters of shape * significant correlations p < 0.0001, Bonferroni-corrected α for six correlation tests = 0.0083.

| Visual Indices | Second Order Spearman Correlation to Perceptual Ratings |
| --- | --- |
| Simple Matching Coefficient | 0.28 * |
| Silhouette | 0.14 * |
| Jaccard Distance | 0.10 * |
| Spatial FFT | 0.02 |
| Binary Images | 0.06 |

### 3.3 Comparison of auditory perceptual ratings to acoustic parameters of pseudowords

Then we compared the auditory perceptual RDM to its within-modal acoustic parameter RDMs (Figure 1, step 3). To determine what auditory stimulus properties may influence the judgment of roundedness or pointedness of pseudowords, we compared the 537×537 RDM for pseudoword judgments with the RDMs for the quantitative indices of these pseudowords. For the following comparisons, we returned to the original set of 537 pseudowords to examine the entire spectrum of rounded/pointedness and the acoustic parameters that may influence these judgments. As an example, Figure 5 compares the dissimilarity matrix of pointedness/roundedness ratings for the pseudowords (left) to that for the temporal FFT: these matrices were positively correlated (Spearman r = 0.28, p < 0.0001).

**Figure 5.**
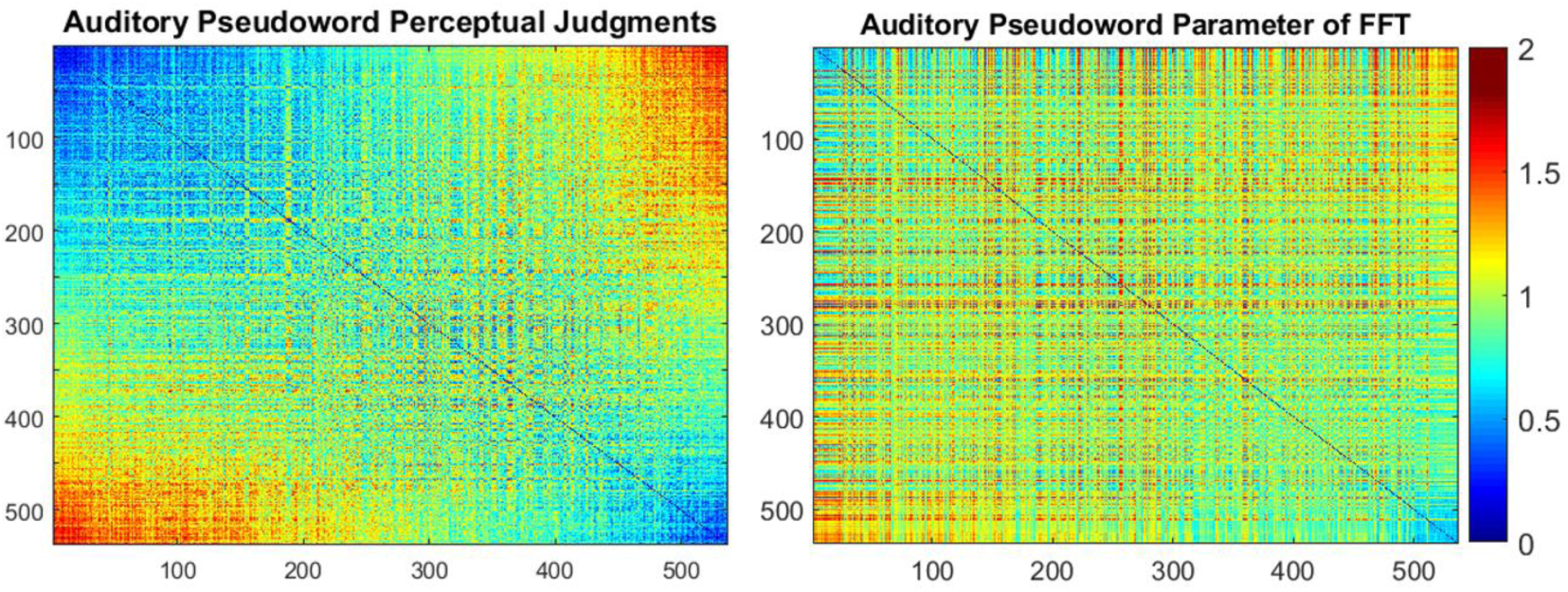
RDMs for the perceptual judgments for pseudowords (left) and one acoustic parameter, the temporal FFT (right). Both RDMs are ordered according to mean perceptual judgments for pseudowords from 1 (rated most pointed) - 537 (rated most rounded). A positive correlation was found between the two RDMs, indicating that the temporal FFT captures acoustic parameters important for auditory perceptual judgments of pointedness/roundedness (Spearman r = 0.28, p < 0.0001).

Table 4 shows that the RDM for pseudoword judgments was positively correlated with the RDMs for shimmer, the temporal FFT (as noted above), mean HNR, speech envelope, and spectral tilt, indicating that these acoustic parameters capture properties that are important for the judgment of roundedness or pointedness for pseudowords. The RDMs for the fraction of unvoiced frames, jitter, mean autocorrelation, period number, pulse number, and pitch standard deviation were not significantly correlated with those for judgments of pseudowords, indicating that these parameters are less important for the judgment of roundedness or pointedness for auditory pseudowords. When comparing perceptual judgments to parameters that comprised a single value per pseudoword using second-order correlations (Methods Table 2), we down-sampled the RDM for the perceptual judgments and binned the acoustic measurements to create 18×18 matrices with 30 pseudowords within each cell. This provided a sufficient number of measures to compute a correlation coefficient that could be entered into each cell of an RDM, while still allowing a sizable enough matrix to be compared with another RDM. Because of the requisite binning procedure for the most highly correlated parameter of shimmer, we instead chose the most highly correlated parameter that was not binned, the temporal FFT, as the example RDM in Figure 5 to illustrate the entire spectrum of pseudoword correlations.

**Table 4.** Correlations between the RDMs for auditory perceptual judgments and quantitative parameters of pseudowords. Significant correlations = * p < 0.001; **p < 0.0001, Bonferroni-corrected α for eleven correlation tests =0.0045.

| Acoustic Parameters | Second Order Correlation to Perceptual Ratings |
| --- | --- |
| Shimmer | 0.30 * |
| Temporal FFT | 0.28 ** |
| Mean HNR | 0.27 * |
| Speech Envelope | 0.14 ** |
| Spectral Tilt | 0.10 ** |
| Fraction of Unvoiced Frames | 0.10 |
| Jitter | -0.01 |
| Mean Autocorrelation | -0.03 |
| Period Number | -0.04 |
| Pulse Number | -0.08 |
| Pitch Standard Deviation | -0.12 |

### 3.4 Comparison of indices of visual and acoustic stimuli

We next compared significant shape and pseudoword parameters across modality (Figure 1, step 4). Because the visual shape and auditory pseudoword judgments involved independent sets of participants, it is possible that participants were making judgments based only on a conceptual representation of rounded or pointed that did not involve sound-symbolic crossmodal associations. If participants were not employing sound-symbolic CCs in their judgments of the pseudowords, we would expect the visual and acoustic parameters that were significant for the two sets of judgments to be unrelated. If instead participants were using a common perceptual framework while judging stimuli from a single modality, particularly the auditory pseudowords, we would expect the auditory and visual parameters to be correlated with one another.

We compared stimulus parameters across modalities in order to determine which of the examined parameters in the two modalities are most likely to be driving sound-symbolic CCs (Figure 6 and Table 5). Once we determined which stimulus parameters were most correlated with the perceptual judgments, we used the RDMs of the two most positively correlated parameters from the visual domain to compare with the two most positively correlated parameters from the auditory domain that would still allow us to maintain a consistent matrix size of 90 by 90. Thus, we chose the simple matching coefficient and silhouette from the visual parameters, and the FFT and speech envelope from the acoustic parameters.

**Figure 6.**
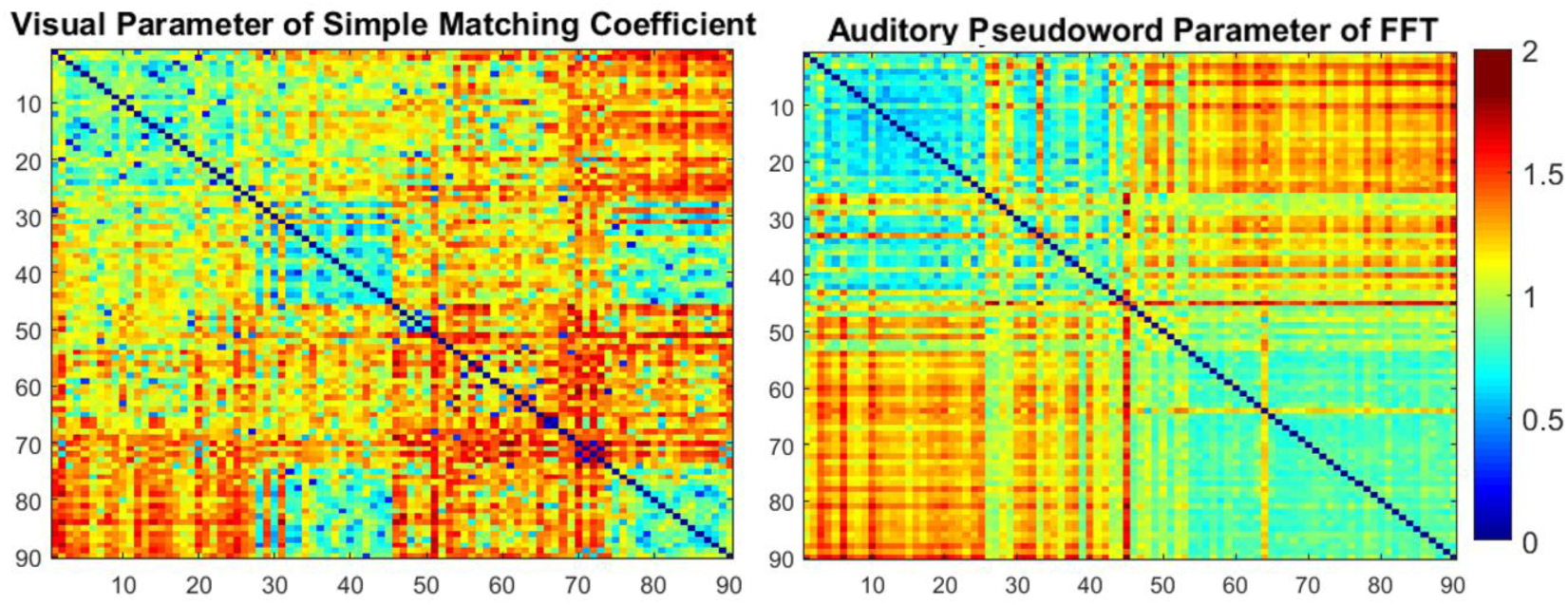
RDMs for the visual simple matching coefficient (left) and the acoustic temporal FFT (right). These RDMs are ordered according to mean perceptual judgments for shapes (left) or pseudowords (right) from 1 (rated most pointed) - 90 (rated most rounded). A positive correlation was found between these RDMs, indicating that the simple matching coefficient for the shapes and the FFT for the pseudowords both relate to properties relevant for the perception of sound-symbolic CCs (r = 0.13, p < 0.0001).

**Table 5.** Correlations between visual and acoustic parameters. Significant correlations = * p < 0.0001, Bonferroni-corrected α for four correlation tests = 0.0125.

| Visual Parameters | Auditory Parameters | Second-Order Spearman<br>Correlation Visual to<br>Auditory Parameters |
| --- | --- | --- |
| Simple Matching Coefficient | Temporal FFT | 0.13* |
| Silhouette | Temporal FFT | 0.03 |
| Simple Matching Coefficient | Speech Envelope | - 0.01 |
| Silhouette | Speech Envelope | 0.02 |

**Figure 7.**
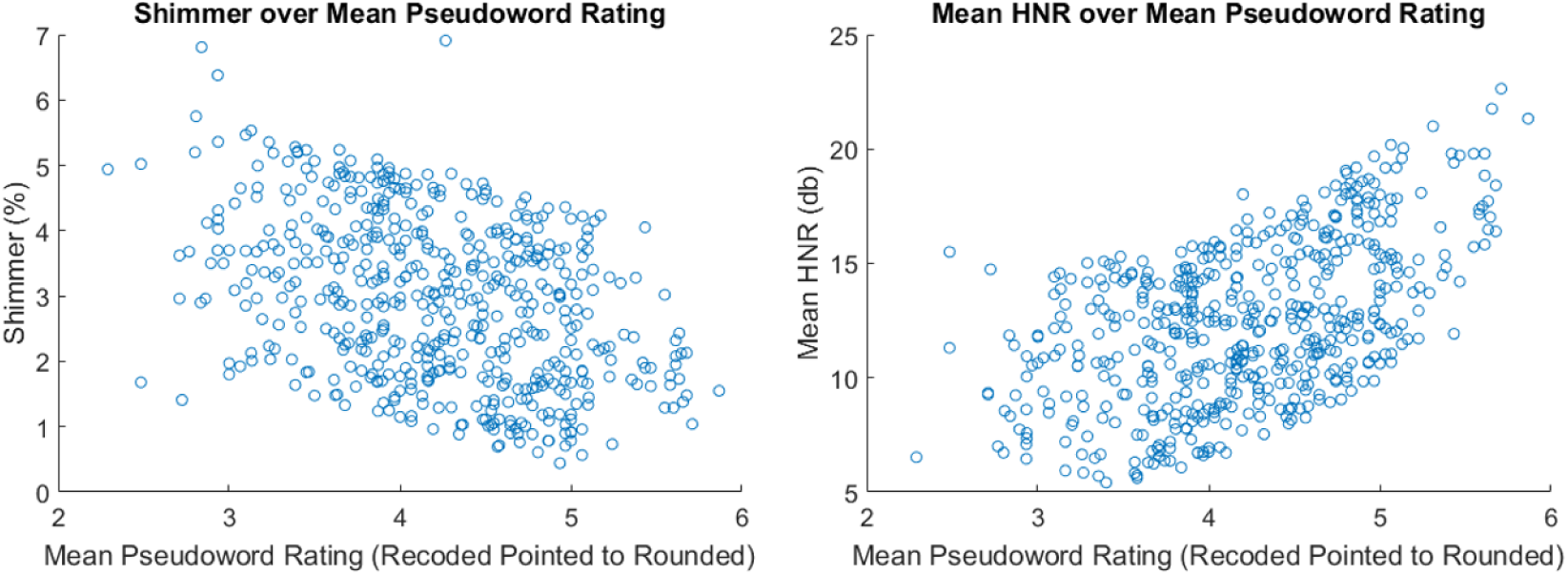
Scatterplots for the acoustic parameters of shimmer (left) and mean HNR (right) over the mean pseudoword ratings recoded across Likert tasks to range from 1-7: pointed to rounded (from left to right on the x axes). Shimmer was negatively correlated, and mean HNR was positively correlated with ratings.

Table 5 shows that the RDM for the simple matching coefficient was significantly correlated with that for the temporal FFT (Spearman r = 0.13, p<0.001: Bonferroni-corrected α for four correlation tests = 0.0125), which suggests that the temporal FFT is an acoustic parameter capturing aspects of the signal that are related to the visual parameters of the shapes indexed by the simple matching coefficient. The silhouette was also weakly positively correlated with the FFT (Spearman r = 0.03, p<0.05) but this correlation did not survive Bonferroni correction. The RDMs for the simple matching coefficient and silhouette were not significantly correlated with that for the speech envelope (all p > .2), indicating that, while the speech envelope is important for auditory judgments of roundedness and pointedness, it may not be directly related to the visual parameters of the simple matching coefficient and silhouette.

### 3.5 Correlations between significant acoustic parameters and auditory perceptual ratings

For the acoustic parameters that yielded RDMs that were correlated with the RDM for the perceptual judgments, we were interested in how these parameters relate to what makes a ‘round’ or a ‘pointed’ pseudoword. Our parameters of shimmer and mean HNR included one measurement per pseudoword, so we were able to take a quantitative approach in correlating shimmer or mean HNR, respectively, with the mean perceptual rating of each pseudoword.

As noted earlier, higher shimmer can indicate increased hoarseness or breathiness in the voice (Teixeira & Fernandes, 2014) and is a measure of amplitude instability in the glottal waveform. We found that shimmer decreased from more pointed to more rounded pseudowords (r = −0.40, p<0.0001) indicating that a pseudoword that is judged as rounded has lower shimmer than one that is judged as more pointed. Mean HNR is a measure of the relative amount of noisiness in the signal with higher values indicating less noise. Thus, an increase in HNR may indicate increased ‘smoothness’ of the waveform (Teixeira & Fernandes, 2014). In contrast to shimmer, there was a positive correlation between mean HNR and ratings from more pointed to more rounded pseudowords (r = 0.53, p<0.0001), indicating that pseudowords that were judged as more rounded had a higher mean HNR than those that were judged as more pointed.

Our parameters of temporal FFT, speech envelope and spectral tilt all included multiple measurements per pseudoword, preventing us from performing a simple correlation between each of these parameters and the pseudoword ratings. Instead, we took a qualitative approach in illustrating how these parameters might vary between a more pointed pseudoword (‘keh-teh’) versus a more rounded pseudoword (‘moh-loh’). For example, the temporal FFT is a recreation of the spectrogram for each pseudoword. Illustrated in Figure 8 (top) is the waveform for ‘keh-teh’ (left) and ‘moh-loh’ (right). The spectrogram (Figure 8, bottom) of the two pseudowords visually illustrates what is captured by the FFT, including the amplitude (degree of dark shading) across time (x-axis) over multiple frequencies (y-axis) for each pseudoword. Figure 8 shows the unvoiced (aperiodic sections of both the spectrogram and the waveform) portion of the pointed pseudoword ‘keh-teh’. Figure 9 illustrates the power spectral density (PSD) of the pointed pseudoword ‘keh-teh’ (left) versus the rounded pseudoword ‘moh-loh’ (right); the slope of the PSD over short time windows illustrates the spectral tilt, another complex acoustic parameter that was important for sound-symbolic judgments of pseudowords. Both Figure 8 and Figure 9 capture the increased power at higher frequencies in the pointed versus the rounded pseudowords.

**Figure 8.**
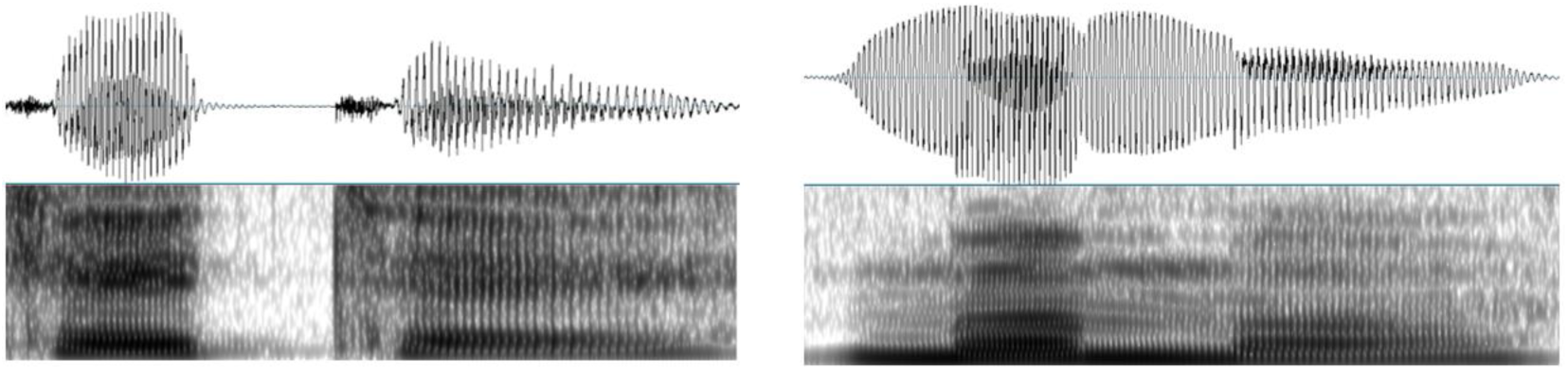
Waveform (top) and spectrogram to illustrate the acoustic parameter of temporal FFT (bottom) for a pointed pseudoword ‘keh-teh’ (left) and a rounded pseudoword ‘moh-loh’ (right). More abrupt changes in the power, especially at higher frequencies, for the pointed pseudoword ‘keh-teh’ versus the rounded pseudoword ‘moh-loh’ were captured by the temporal FFT.

**Figure 9.**
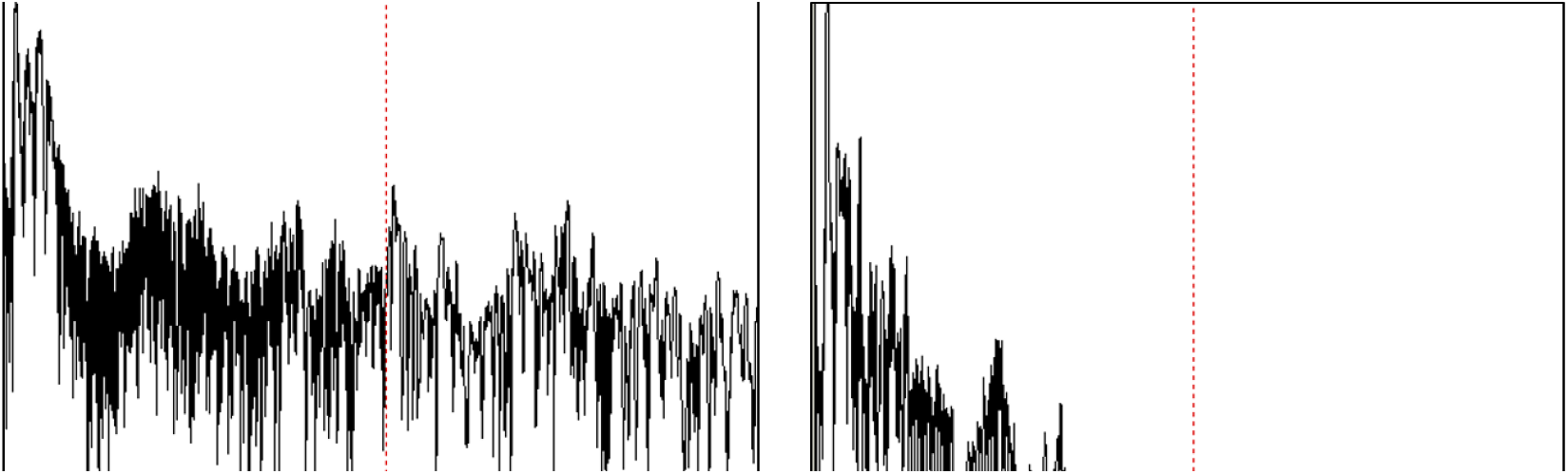
PSD showing power across time for two pseudowords illustrates the acoustic parameter of spectral tilt for a pointed pseudoword ‘keh-teh’ (left) and a rounded pseudoword ‘moh-loh’ (right). The PSD and spectral tilt capture increased power at higher frequencies for the pointed pseudoword ‘keh-teh’ versus the rounded pseudoword ‘moh-loh’.

Finally, we qualitatively examined the speech envelope for a pointed pseudoword (‘keh-teh’, Figure 10, left) versus a rounded pseudoword (‘moh-loh’, Figure 10, right). The speech envelope encapsulates the shape of the waveform in continuous measurements of amplitude across time. This measurement is perhaps the most intuitive illustration of the ‘shape’ of a pseudoword and how the jagged shape of a pointed pseudoword (left) differs from the rolling, full shape of a rounded pseudoword (right).

**Figure 10.**
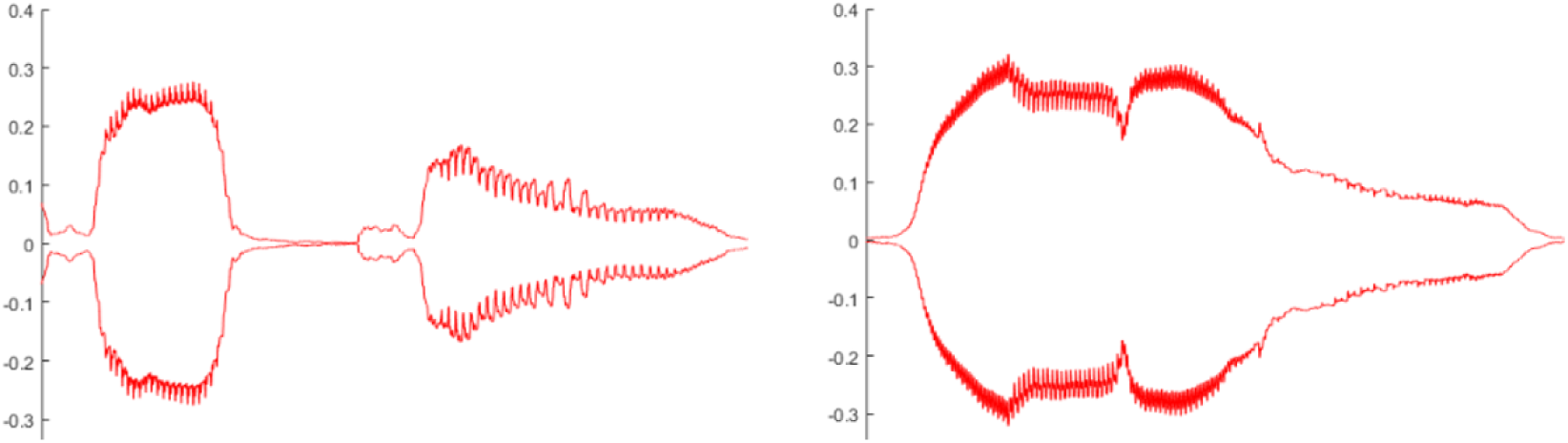
The acoustic parameter of speech envelope for a pointed pseudoword ‘keh-teh’ (left) and a rounded pseudoword ‘moh-loh’ (right) illustrates the “shape of sound” as measured by the amplitude across time.

## 4. DISCUSSION

### 4.1 Sound-symbolic crossmodal correspondences and perception

The present study is the first to systematically examine both the acoustic and the visual drivers of sound symbolism in the sound-shape domain within the same experimental paradigm and using RSA; previous studies have only examined such parameters separately (visual: Chen et al., 2016; acoustic: Knoeferle et al., 2017). Additionally, our stimulus sets comprising 90 shapes and 537 pseudowords are more extensive than in previous studies (e.g. Knoeferle et al., 2017 included 10 shapes and 100 pseudowords). This advantage allowed us to explore a spectrum of rounded to pointed shapes and pseudowords, with significant power. Our study is also the first to investigate the similarities in perceptual judgments of rounded/pointedness for shapes compared to pseudowords and to further use these similarities to determine which visual and acoustic parameters are important for sound-symbolic CCs both within and across modalities. Despite using different parameters to make sound-symbolic judgments, raters were able to use a common scale of roundedness/pointedness across the two different modalities, as indicated by the similarity in the RDMs for shape versus pseudoword judgments. This is not necessarily surprising, given that participants were asked to use the same scales in both modalities, but the fact that different groups of participants, using two opposing scales in each modality, produced ratings whose dissimilarity matrices were highly correlated across modalities, is evidence that the sound-symbolic CC studied here is robust. The strong crossmodal correlation provides a firm basis for further exploration of the principal aim of the present study, characterizing stimulus parameters that may underpin sound symbolism.

For the shapes, the parameters of simple matching coefficient, silhouette, and Jaccard distance were predictive of individuals’ judgments of visual roundedness/pointedness, as revealed by correlations between the RDMs for these parameters and the perceptual ratings. For the pseudowords, RDMs showed that the parameters of shimmer, temporal FFT, mean HNR, speech envelope, and spectral tilt were predictive of individuals’ judgments of the roundedness/pointedness of sounds. As further illustrated by inspecting the spectrograms for the temporal FFT and spectral tilt for pseudowords judged as very pointed and very rounded, increased power in higher frequencies led to perceptual ratings that were more pointed. Inspecting the corresponding speech envelopes indicated that smoother and more gradual changes in amplitude were associated with perceptual ratings that were more rounded. Examination of correlations revealed that the acoustic parameter of shimmer, which can indicate increased breathiness (Teixeira & Fernandes, 2014) and amplitude variation and instability, *decreased* progressively as the ratings of roundedness increased, whereas the acoustic parameter of mean HNR, which indicates the ratio of periodic or smooth to aperiodic or noisy parts of a vocal signal, *increased* progressively as the ratings of roundedness increased.

Using RSA to compare across acoustic and visual parameters, we were able to determine that the spectral composition of auditory pseudowords as measured by the temporal FFT may be employed by listeners in a similar way as the visual index of shape quantified by the simple matching coefficient. This evidence suggests that even for unisensory stimuli, a parameter of sound in the case of pseudowords can be perceived as equivalent to a parameter of vision in the case of shapes, offering a basis for the sound-symbolic pseudoword-shape CC.

Our findings extend previous studies on sound symbolism suggesting that sound-symbolic ratings are grounded in broad categorical contrasts, such as the phonological features characterizing obstruent (e.g. ‘p’, ‘t’, and ‘k’ sounds in English) or sonorant (e.g. ‘m’, ‘n’, or ‘l’ sounds in English) consonants (Nielsen & Rendall, 2011; Fort, Martin, & Peperkamp, 2014) or the presence or absence of corners in a visual shape (Chen et al., 2016). In addition, individuals seem to base sound-symbolic CCs on intrinsic physical properties of shapes and speech sounds.

We also provide nuance to the assumption that the sound-symbolic judgments of pseudowords are based on a conception of rounded and pointedness that derive from orthography of the pseudowords (Cuskley et al., 2015) or in the shape of the lips when producing certain sounds (Ramachandran & Hubbard, 2001; Namy & Nygaard, 2008). Sound-symbolic CCs are not just supported by a conception of shape that links the visual to the auditory domain through production of the sounds or the shape of their written form. Our findings show that sound-symbolic CCs are also related to fundamental physical properties of the visual and auditory stimuli that evoke such CCs.

Visual and acoustic parameters of shapes and pseudowords are of course closely interwoven with the visual and phonetic categories into which the shapes and pseudowords can be divided (Nielsen & Rendall, 2011). Vowel frontness and lip rounding are associated with the frequencies of the first and second formants. Consistent with Nielsen and Rendall’s (2011; see also Fort, Martin, & Peperkamp, 2014) finding that lip rounding and vowel frontness are important predictors of judgments of roundedness in pseudowords, Knoeferle et al. (2017) found that the relative frequency of the second formant was an important predictor of judging both the size and the roundedness of shapes. We have evidence that supports Knoeferle et al. (2017) in that the physical parameters of sound are particularly vital to the judgment of roundedness/pointedness of pseudowords, especially from our measurement parameters like the temporal FFT that capture multiple acoustic dimensions of a speech sound. In our results, energy in higher frequencies in the temporal FFT were associated with pointedness, and consistent with the formant analyses of Knoeferle et al. (2017), energy in lower frequencies were associated with roundedness. However, to the extent that physical properties and phonetic features and visual categories are linked, we cannot at present resolve the question of whether sound-symbolic CCs originate from low-level acoustic and visual properties or whether they stem from more general features of phonetic categories.

Our study represents an initial foray into testing the role of multiple visual and acoustic parameters in sound-symbolic CCs. However, our investigation of visual and acoustic parameters was not intended to be exhaustive, and there may be other visual and acoustic parameters not explored here that are important for the perception of sound-symbolic CCs. In particular, we did not find visual or acoustic parameters whose RDMs were significantly negatively correlated with the RDMs of the corresponding perceptual judgments. However, we did find that shimmer, which indicates increased hoarseness or breathiness in the voice (Teixeira & Fernandes, 2014) was negatively correlated with ratings of roundedness. In contrast, the mean HNR was positively correlated with roundedness ratings, and mean HNR is an index of noisiness. Higher HNR indicates less noise and was correlated with higher ratings of roundedness, which is consistent with the fact that lower mean HNR indicates increased hoarseness in the voice (Teixeira & Fernandes, 2014).

In addition, our visual shapes were simplistically constructed to have identical grayscale contrasts and lacked internal visual patterns, which limited the complexity of visual properties that we could sensibly investigate, compared to others with more variable sets of images (Jonas, Spiller, & Hibbard, 2017). The simplicity of our shapes may explain why spatial FFT was not a significant contributor to the perceptual judgments of shape (Petrou & Petrou, 2011). Future studies should examine complex images or examine visual properties important for other types of CCs (Chen et al., 2016; Jonas, Spiller, & Hibbard, 2017).

It is interesting that acoustic properties associated with variation in voice quality, such as shimmer and mean HNR (Teixeira & Fernandes, 2014), were positively correlated with the perceptual judgments. One possible explanation is that the speaker who recorded the stimuli produced pseudowords that were expected to be perceived as rounded or pointed differently, implicitly varying these acoustic properties (Marcel, 1983). However, shimmer and mean HNR reflect properties of speech that are less likely to be affected by bias, as opposed to a more easily altered parameter such as pitch, so the explanation of an unconscious bias is unlikely to completely account for the correlations of these acoustic properties with sound-symbolic judgments. Prosodic cues that speakers are able to modulate can impart meaning of perceptual details in the visual modality, such as color brightness (Tzeng et al., 2018). Sound-symbolic CCs may similarly be affected by speakers’ intentional or unintentional changes in vocal patterns to impart meaning (Parise & Pavani, 2011). Future experiments that intentionally vary acoustic properties such as shimmer, mean HNR, and speech envelope within an extensive stimulus set may shed more light on the relative influence of these acoustic properties on sound-symbolic CCs.

Neuroimaging studies of sound-symbolic CCs may further elucidate the nature of the relationship between sound and meaning in these types of stimuli. Currently, there are few published neuroimaging studies in the area of sound symbolism. Perhaps in the presence of sound-symbolic CCs, individuals engage areas of the brain linked to the perception of speech and multisensory integration, like the posterior superior temporal sulcus (Beauchamp, Nath, & Pasalar, 2010). Alternatively, perhaps the presence of sound-symbolic CCs would differentially engage participants’ frontoparietal areas that are linked to attention (McCormick et al., 2018; Peiffer-Smadja & Cohen, 2019) and abstract categorization (Revill et al., 2014). Revill et al. (2014) showed that left superior parietal lobule exhibited heightened activation during presentation of sound-symbolic versus non-symbolic words, while fractional anisotropy in the superior longitudinal fasciculus, an area implicated in phonological processing and multisensory facilitation (Brang et al., 2013), correlated with performance of a sound-symbolic task. These authors interpreted their findings as supporting the idea that sound symbolism is processed in areas important for multisensory integration.

### 4.2 Sound symbolism in language acquisition

Neuroimaging methods are one type of approach to the ubiquitous question of why sound symbolic CCs exist in natural language and how they arise (Sidhu & Pexman, 2017). One explanation to this question is that these associations are a necessary part of language acquisition (Imai & Kita, 2014). Researchers investigating the role of sound symbolism have found evidence to suggest that sound-symbolic CCs are used to determine meanings of words based on individual properties of words and on the broader category to which a word belongs. On the one hand, sound-symbolic CCs provide a tool for children to acquire language by learning the meaning of individual words. Imai et al. (2015) found that the sound structure of individual words bolstered children’s abilities to learn the meaning of that word (see also Ozturk, et al., 2013; Tzeng, Nygaard, & Namy, 2017). This theory is known as the bootstrapping hypothesis (Imai & Kita, 2014). The evidence for this theory indicates that sound-symbolic CCs could be based in individual characteristics of words, at least for young children. Gasser (2004) used computer simulations to illustrate the advantage of sound symbolism in small vocabularies, and this advantage was supported by Brand et al.’s (2018) study of adults learning the individual meanings of smaller sets of pseudowords.

Studies of sound-symbolic CCs also show that they are important for determining meaning across general categories. The findings from Gasser (2004) also indicate that the advantage of determining the specific meanings of words using sound symbolism is diminished as the vocabulary increases (although see Revill et al., 2018). Whether sound symbolism facilitates learning categories or specific word meanings in adults, the idea that sound symbolism can benefit adult learners has been recently validated with eye-tracking and other behavioral studies (Brand et al., 2018; Revill et al., 2018).

Regardless of whether or not sound-symbolic CCs originate from aligned percepts across sensory modalities, having a motivated connection between words and meaning has been suggested to facilitate language learning. The pervasiveness of sound symbolism across languages supports the theory that it may have an important role in language learning (Imai et al., 2015; Kantartzis, Imai, & Kita S, 2011; Tzeng, Nygaard, & Namy, 2017). Individuals are even able to correctly assign meanings of synonym/antonym pairs above chance for languages with which they are unfamiliar (Kunihara, 1971; Nygaard et al., 2009; Tzeng, Nygaard, & Namy, 2016). Our study suggests that one potential motivated connection between sound and meaning is the set of acoustic properties to which listeners are sensitive and that translate systematically into a set of visual properties. If we consider the case of young children learning words, this set of acoustic properties should be connected crossmodally to a set of visual properties in order for the bootstrapping hypothesis to be important for concrete meanings such as shape or size. Sound and meaning may be mapped not only at the word or phonetic level, but perhaps also at the more finely tuned level of acoustic properties and in the case of concrete, physical meanings, also at the level of visual properties. It is also interesting that iconic words such as “moo” and “splash” lead to better performance than non-iconic words on reading aloud and auditory lexical decision tasks in aphasic individuals (Meteyard et al., 2015).

### 4.3 Theoretical accounts of sound symbolic correspondence

An active question in the field of sound symbolism that may serve to explain why these associations exist has been whether sound symbolism is based on a gradation or spectrum of sounds and meanings (e.g. a range of sounds to symbolize the spectrum of brightness from dim to medium to bright) or whether it is a matter of relative opposites (e.g. contrasting phonetic categories to symbolize dim versus bright). This question is known as the continuous-contrastive marking problem (Thompson & Estes, 2011) and is central to the theories on the development of sound symbolism in natural language. The statistical theory states that sound symbolism became incorporated into linguistic reference as languages evolved because listeners consistently registered the random distribution of sounds in language. Certain sounds became statistically more likely to be present in words with specific meanings by chance, such as the phonological differences between English nouns and verbs (Farmer, Christiansen, & Monaghan, 2006). This theory most readily explains contrastive sound-to-meaning mappings in that the pairing between a sound and its meaning is initially arbitrary but over time becomes symbolic because of the statistical co-occurrence of those sounds (Thompson & Estes, 2011).

In contrast, the crossmodal theory states that sound symbolic mapping in language was spurred by listeners matching gestures and acoustic properties of speech to physical properties of stimuli in vision or other sensory modalities (Ramachandran et al., 2001). Gestures, acoustic properties of speech sounds, and physical properties in vision or other sensory modalities can all be graded and finely tuned to portray a continuous spectrum of meaning. For example, Thompson and Estes (2011) found that the number of large-sounding phonemes in an object label was associated with object size and dimensions. Knoeferle et al. (2017) extended these findings in support of the crossmodal theory by expanding from sound-size to sound-shape symbolism and examined acoustic properties of vowels. They found that sound-symbolic crossmodal judgments varied linearly as a function of the second and third formants and that different acoustic properties were associated with the two different meaning dimensions. Future studies will need to explore the acoustic and visual parameters discussed here, among others, in the context of multiple dimensions of sound-symbolic meaning to determine which parameters, if any, may be important for sound symbolism in general versus for a specific dimension of meaning.

We here provide a key link between continuous acoustic properties and continuous visual properties and show that the perception of roundedness and pointedness along a spectrum in the visual and auditory domains is correlated with similar variations along a spectrum in the simple matching coefficient for visual shapes and the FFT for auditory pseudowords. This finding supports the crossmodal theory that sound symbolism resulted from links across a continuous spectrum for certain auditory and visual properties.

Future work could extend the current study by exploring additional CCs, such as mappings between sound and size (Jonas, Spiller, & Hibbard, 2017), taste (Wang et al., 2016) or emotion (Aryani et al., 2018). It would also be of interest to investigate whether children of different ages or elderly adults are sensitive to similar acoustic and visual properties or if sound-symbolic CCs correspondences change across the lifespan. Studies such as these would further illuminate the potential underlying perceptual and neural mechanisms of sound-symbolic CCs.

## 5. CONCLUSION

In sum, the current findings suggest that sound-symbolic CCs are driven by distinct acoustic and visual parameters. Specifically, pairwise dissimilarities between the simple matching coefficient, silhouette, and Jaccard distance were correlated with pairwise dissimilarities between judgments of the roundedness/pointedness of visual shapes, while pairwise dissimilarities between the shimmer, FFT, mean HNR, speech envelope, and spectral tilt were correlated with pairwise dissimilarities between sound-symbolic judgments of the roundedness/pointedness of auditory pseudowords. Individuals are not only sensitive to sound-symbolic CCs but are also able to employ visual associations (i.e., rounded vs. pointed) for pseudowords even in the absence of the physical shapes that the sounds may relate to. These findings suggest that sound symbolism is not only a set of visual categorical contrasts that are instantiated by phonetic properties of words, but that individuals are also able to base their sound-symbolic judgments on a continuum of basic visual and auditory properties. Taken together, these findings imply that the relationship between words and their meanings can have a non-arbitrary basis at the level of auditory and visual stimulus properties, and highlight the importance of sound symbolism in natural language.

## ACKNOWLEDGEMENTS

This work was supported by grants to KS and LN from the National Eye Institute at the NIH (R01EY025978) and the Emory University Research Council. Support to KS from the Veterans Administration and to SML from the Laney Graduate School is also acknowledged. We thank Jee Young Kim and Valentin Lazar for their advice and assistance.

## Author contributions

S.M.L., K.R.M., S.L., K.S. and L.C.N. designed research; K.R.M. created all stimuli; S.M.L. and K.R.M. performed research; S.M.L. analyzed data; and S.M.L, S.L., K.S. and L.C.N. wrote the paper.

